# PRIME: A modular plasmid-based platform for continuous in vivo evolution across bacteria

**DOI:** 10.64898/2026.09.28.754915

**Authors:** Erik Dechant, Lena K. Träger, Srdan Sikanjic, Lavanja Varathan, Maike Otto, Nicolas Huguenin-Dezot

## Abstract

Directed evolution enables the engineering of proteins with novel or improved functions, yet existing approaches often require extensive manual intervention and are difficult to sustain over long evolutionary trajectories. Here we introduce PRIME (<u>P</u>rotein-primed <u>R</u>eplication for *In vivo* <u>M</u>utagenesis and <u>E</u>volution), a plasmid-based orthogonal DNA replication system for continuous directed evolution in *Escherichia coli*. PRIME harnesses protein-primed DNA replication to establish an autonomous replicon that operates independently of the host genome. Coupling this system to an error-prone DNA polymerase enables targeted diversification of constructs encoded on the orthogonal plasmid while preserving genomic integrity. Using only standard laboratory equipment, we demonstrate the evolution of a construct expressing msfGFP with an 11.8-fold increase in fluorescence. The platform is fully compatible with standard molecular biology workflows and requires no specialised instrumentation. We further demonstrate its portability across bacterial hosts by establishing PRIME in *Pseudomonas putida*. PRIME thus provides a broadly accessible framework for continuous *in vivo* evolution, with wide-ranging applications in protein engineering and synthetic biology.

## Introduction

Proteins are involved in all processes of life and play crucial roles in endowing living matter with structure and function. The immense array of protein shapes and functions is the result of very long evolutionary processes of random mutations and selection, that let emerge successful versions of each protein for the different requirements of life. Directed evolution is a technique that aims to replicate and speed up this evolutionary process in the laboratory by introducing mutations and providing targeted selection conditions^1^. Historically, mutations were introduced into genes by several *in vitro* techniques and mutants with improved or novel functions were subsequently selected. However, these techniques are labour-intensive and low throughput, and therefore only allow short mutational paths to be traversed. An ideal system would allow for mutants to be generated, and simultaneously selected for *in vivo,* without manual intervention, allowing the evaluation of large numbers of mutations and hence, cover longer mutational paths. Importantly, the mutagenesis needs to be specific for the gene(s) of interest to avoid mutations that would affect the genome of the host in a detrimental way and/or perturb the selection process.

An effective strategy to achieve this is to restrict mutagenesis to a DNA episome (i.e., a plasmid) whose replication is independent of the host replication machinery and instead relies on its own dedicated replication enzymes. Because such a plasmid replicates orthogonally to the host genome, the use of an error-prone DNA polymerase allows mutations to be introduced selectively into the episomal DNA without accumulating deleterious mutations in the host genome. At the same time, generating mutations directly *in vivo* greatly increases the throughput of directed evolution experiments, eliminating the laborious and time-consuming steps associated with traditional *in vitro* approaches. The advantages of using an orthogonal DNA replication system for continuous directed evolution in a living organism has been demonstrated in 2014 in a landmark paper published by the group of Chang Liu^2^. Since its publication, this system has been used successfully to evolve numerous proteins through long adaptive mutational pathways^3,4^. Several years later, systems working in *Bacillus thuringiensis*^5^ and *Escherichia coli*^6–8^ were also described, both taking advantage of DNA replication systems from bacteriophages. Our laboratory has specialised in the study of the bacteriophage PRD1 bacteriophage and its protein-primed DNA replication machinery. Building on this expertise, we sought to develop an orthogonal DNA replication platform based on the PRD1 replication system. Previous work had shown that the PRD1 DNA replication machinery can be used to establish orthogonal replication in *E. coli*^6^. However, several groups including ours, have observed toxic effects when expressing PRD1 replication proteins in *E. coli*^9–12^. These limitations have so far prevented the stable cloning of plasmids containing the full replication machinery and hindered the development of a fully plasmid-based orthogonal replication system based on this organism.

Here, we present PRIME (<u>P</u>rotein-primed <u>R</u>eplication for *In vivo* <u>M</u>utagenesis and <u>E</u>volution), a system based on the PRD1 DNA replication machinery that enables the establishment of orthogonal DNA plasmids in *E. coli*. The DNA replication machinery is encoded on a single plasmid (pORM, for <u>O</u>rthogonal <u>R</u>eplication <u>M</u>achinery), while two accessory plasmids facilitate transformation (pOTH, for <u>O</u>rthogonal <u>T</u>ransformation <u>H</u>elper) and production of linear orthogonal replicons (pORFITR, for <u>O</u>rthogonal <u>R</u>eplicons <u>F</u>lanked by Inverted <u>T</u>erminal <u>R</u>epeats) (**Fig. 1**). PRIME readily integrates into standard molecular biology workflows and does not require specialised equipment, making it accessible to a broad range of laboratories.

**Fig. 1:**
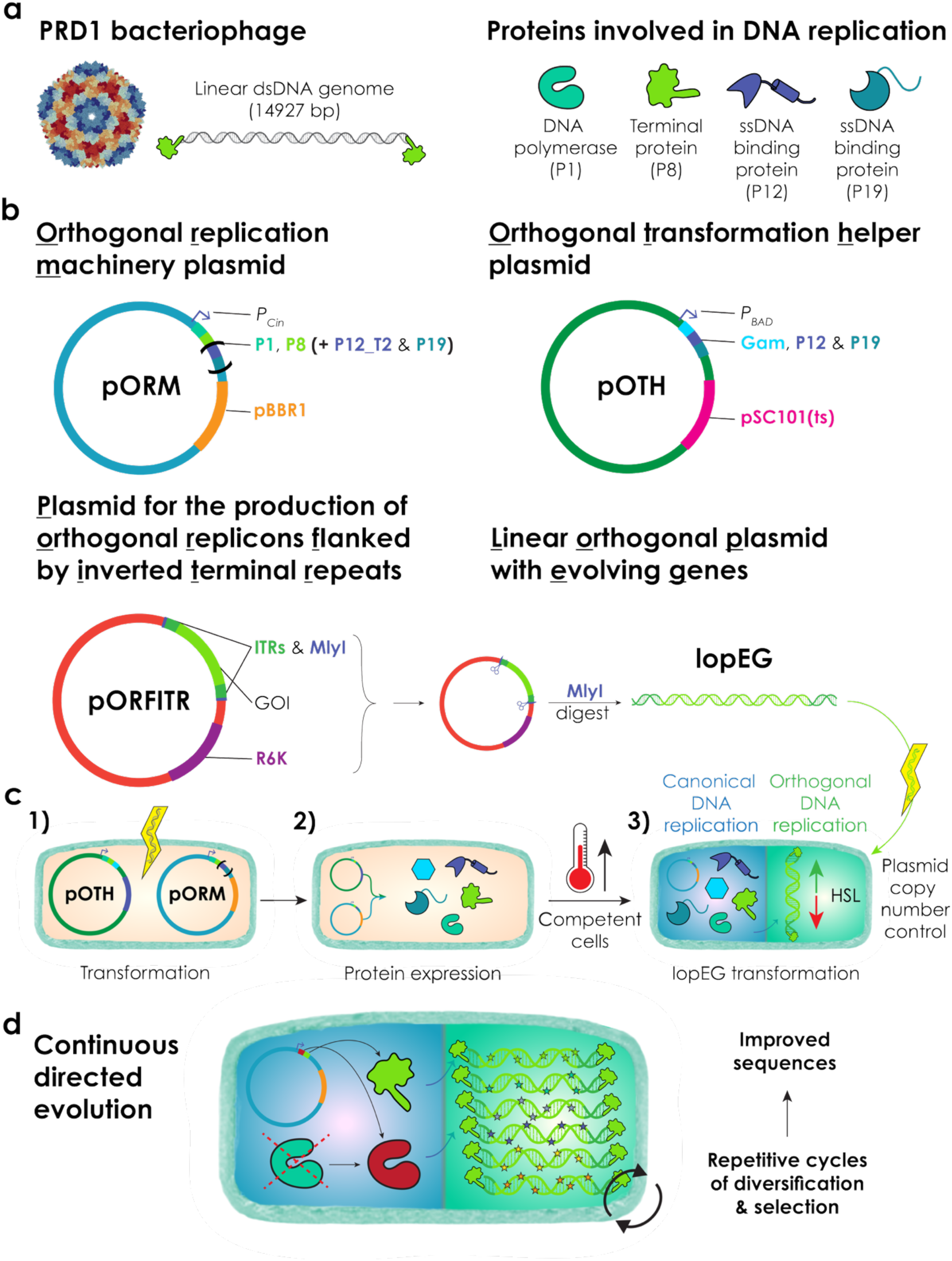
The PRIME system (<u>P</u>rotein-primed <u>R</u>eplication for *In vivo* <u>M</u>utagenesis and <u>E</u>volution). **(a)** The PRD1 bacteriophage possesses a 14927 bp genome encoding four proteins involved in protein-primed DNA replication: the DNA polymerase P1, the terminal protein P8, and the single-stranded DNA-binding proteins P12 and P19. **(b)** PRIME comprises four plasmids: pORM (<u>O</u>rthogonal <u>R</u>eplication <u>M</u>achinery), pOTH (<u>O</u>rthogonal <u>T</u>ransformation <u>H</u>elper), pORFITR (<u>O</u>rthogonal <u>R</u>eplicons <u>F</u>lanked by Inverted <u>T</u>erminal <u>R</u>epeats), and lopEG (<u>L</u>inear <u>O</u>rthogonal <u>P</u>lasmid with <u>E</u>volving <u>G</u>enes). The pORM plasmid carries the genes required for protein-primed DNA replication under control of the inducible *P_Cin_* promoter. P12_T2 and P19 are indicated in brackets, as these proteins are dispensable for orthogonal plasmid replication but are present in the pORM_P12_T2 construct. pORM additionally contains the broad-host-range pBBR1 origin of replication, enabling propagation in a wide range of bacterial hosts. The pOTH plasmid encodes accessory proteins facilitating lopEG transformation. Owing to their toxicity, these proteins are expressed under control of the inducible *P_BAD_* promoter. The temperature-sensitive pSC101(ts) origin enables efficient curing of pOTH following lopEG establishment. The pORFITR plasmid facilitates cloning and production of lopEG plasmids. It contains two inverted terminal repeat (ITR) regions flanked by MlyI recognition sites. As a Type IIS restriction enzyme, MlyI cleaves outside of its recognition sequence, enabling blunt cleavage precisely at the edges of the ITRs. The R6K conditional origin of replication prevents plasmid propagation in cells lacking the Pir protein, allowing direct transformation of lopEG plasmids without gel purification. **(c)** Workflow for establishment of orthogonal plasmids in *E. coli*. (1) Cells are first transformed with pORM and pOTH. (2) Expression of the replication proteins is induced, and the temperature is increased prior to lopEG transformation to inhibit further replication of pOTH. (3) lopEG plasmids are electroporated into competent cells expressing the PRD1 replication machinery. Orthogonal plasmid copy number can subsequently be tuned using the OHC14-HSL inducer. **(d)** PRIME enables continuous directed evolution experiments. Replacement of the wild-type P1 polymerase (green) with an error-prone variant (red) promotes the generation of DNA libraries directly *in vivo* without the need for repeated manual intervention. Iterative cycles of diversification and selection subsequently enable the rapid evolution of constructs with improved properties.

We further show that PRIME can be established in an additional biotechnologically relevant bacterium, such as *Pseudomonas putida*. We demonstrate that we can control the orthogonal plasmid copy number in both *E. coli* and *P. putida* by using the OHC14-HSL (HSL)^13^ inducer.

Finally, using a simple experimental workflow, we show that PRIME can rapidly generate improved constructs in *E. coli*, increasing the fluorescence intensity per cell of strains expressing monomeric superfolder green fluorescent protein (msfGFP) 11.8 fold in 7 rounds of selection. Together, the different workflows and plasmids presented here aim to broaden access to directed evolution technologies beyond specialist laboratories.

## Results

### A plasmid-based system for orthogonal plasmid transformation

The genome of the PRD1 bacteriophage is 14927 bp in length and encodes four proteins which have been identified as being involved in DNA replication: P1 (DNA polyermase^14^), P8 (terminal protein^15^), P12 (single-stranded DNA binding protein^9,16^) and P19 (single-stranded DNA binding protein^17^) (**Fig. 1a**). Attempts to establish a plasmid carrying these genes, both by in-house cloning and through commercial gene synthesis providers, were initially hindered by recurrent cloning failures, most likely caused by the toxicity of the P12 protein^9^. To overcome this limitation, we designed a modified operon in which P12 was replaced by a truncated variant (P12_T2) that we had previously shown to retain DNA-binding activity while displaying reduced cellular toxicity^9^. In addition, P1, P8 and P19 were codon-optimised, aiming to disrupt putative internal promoter elements within the original operons, which may have contributed to the observed toxicity. This redesigned architecture enabled the successful construction of a stable plasmid that was tolerated by host cells (**Fig. 1b**). The genes encoding P1, P8, P12_T2, and P19 were furthermore placed under the control of the *P_Cin_* promoter^13^, allowing tuneable expression through modulation of HSL concentration. We also selected the pBBR1 origin of replication for the construction of this plasmid because of its broad host range, including its compatibility with both *E. coli* and *P. putida*. This construct was designated the <u>O</u>rthogonal <u>R</u>eplication <u>M</u>achinery plasmid (pORM).

In addition to pORM, we built another plasmid aiming at transiently supplying proteins stabilising the naked linear DNA fragments after the transformation. We named this plasmid <u>O</u>rthogonal <u>T</u>ransformation <u>H</u>elper plasmid (pOTH) (**Fig. 1b**). This plasmid contains an operon with the native sequences of P12 and P19 from the PRD1 bacteriophage and the Gam protein from the lambda bacteriophage, which have been shown to help with the transformation of linear double-stranded DNA fragments^6^. The expression of these proteins is tightly regulated by the presence of an arabinose promoter upstream of the gene cassette. To efficiently remove pOTH after transformation, we included a temperature sensitive pSC101 ori (pSC101 ts), which allow the curing of the plasmid by growing cells at 37 °C^18^.

We next constructed a plasmid that enables the facile generation of linear DNA fragments flanked by inverted terminal repeats (ITRs). This plasmid was named plasmid for the production of <u>O</u>rthogonal <u>R</u>eplicons <u>F</u>lanked by Inverted <u>T</u>erminal <u>R</u>epeats (pORFITR) (**Fig. 1b**). Using this plasmid eliminates the need for polymerase chain reaction (PCR)-based assembly of linear DNA fragments, which is often challenging due to the substantial length (110 bp) of the PRD1 inverted terminal repeats (ITRs) and their propensity to form secondary structures^19^. To facilitate cloning between the ITR regions, two restriction sites (FseI and SbfI) were introduced at each ITR end. In addition, MlyI recognition sites were positioned flanking the ITRs, enabling precise blunt-end cleavage at the edge of each ITR and direct excision of the fragment required for subsequent transformation. To prevent plasmid recircularisation and undesired circular plasmid propagation following transformation, pORFITR incorporates an R6K conditional origin of replication, thereby restricting replication to *pir*-expressing strains^20^. As a result, the plasmid can be digested with MlyI, dephosphorylated and directly transformed after reaction clean-up. The resulting linear orthogonal plasmid resulting from the digestion of the pORFITR plasmid was designated <u>L</u>inear <u>O</u>rthogonal <u>P</u>lasmid with <u>E</u>volving <u>G</u>enes (lopEG) (**Fig. 1b**).

Using this set of plasmids, we established a streamlined workflow for the efficient generation and transformation of orthogonal replicons in *E. coli* (**Fig. 1c**). In brief, cells are first co-transformed with the pOTH and pORM plasmids (**Fig. 1c[1]**). Competent cells are subsequently prepared while expressing the PRD1 replication machinery from the pORM plasmid together with accessory proteins facilitating transformation from the pOTH plasmid (**Fig. 1c[2]**). Shortly before lopEG transformation, the temperature of the competent cell culture is raised to 37 °C to initiate curing of the pOTH plasmid (**Fig. 1c[3]**). After transformation, the copy number of the lopEG plasmid can be finely tuned by adjusting the concentration of HSL. Replacing the wild-type PRD1 DNA polymerase with an error-prone variant on pORM allows continuous directed evolution to be performed following this transformation protocol (**Fig. 1d**).

### P12 supports lopEG transformation but is not required for plasmid maintenance

While working with pORM_P12_T2 (**Fig. 1b**), we observed that this plasmid occasionally accumulated frameshifts and non-sense mutations disrupting the P12_T2 protein. This prompted us to investigate whether P12_T2 on pORM was required in addition to the P12 protein provided by the pOTH plasmid to perform lopEG transformation. To test this, we constructed a minimal version of the pORM plasmid containing only P1 and P8 (named pORM hereafter) (**Fig. 1b**). Using this minimal version of the plasmid, we repeated our standard transformation protocol (**Fig. 1c**) and found that cells were able to replicate lopEG in the presence of pORM and absence of pOTH, demonstrating that P12 is dispensable for PRD1 DNA replication. This minimal architecture also circumvents the deleterious effects associated with P12 expression that we previously observed^9^. From these results, we conclude that P12_T2 supports the transformation^6^ of lopEG but is dispensable for its subsequent replication and maintenance.

### P8-capped lopEG plasmids can be purified and transformed into cells expressing only P1 and P8

To further investigate whether P12 and P19 were required after the initial transformation step, we purified lopEG plasmids from previously transformed cells and electroporated them into cells expressing only P1 and P8 from pORM, in the absence of pOTH. This experiment yielded transformants, demonstrating that P12 and P19 are dispensable for the transformation, replication and maintenance of lopEG once the DNA is already P8-capped. Together with previous work showing that P12 is required for efficient transformation of linear PRD1-derived DNA^6^, these results indicate that the requirement for P12 is specific to the initial introduction of uncapped DNA rather than subsequent replication and maintenance of the P8-capped replicon.

### lopEG can be transformed in *P. putida* bearing pORM

Building on our observation that lopEG plasmids capped by P8 can be efficiently transformed into *E. coli* cells expressing only pORM, we tested whether our transformation protocol could be adapted to additional biotechnologically relevant bacteria (**Fig. 2**).

**Fig. 2:**
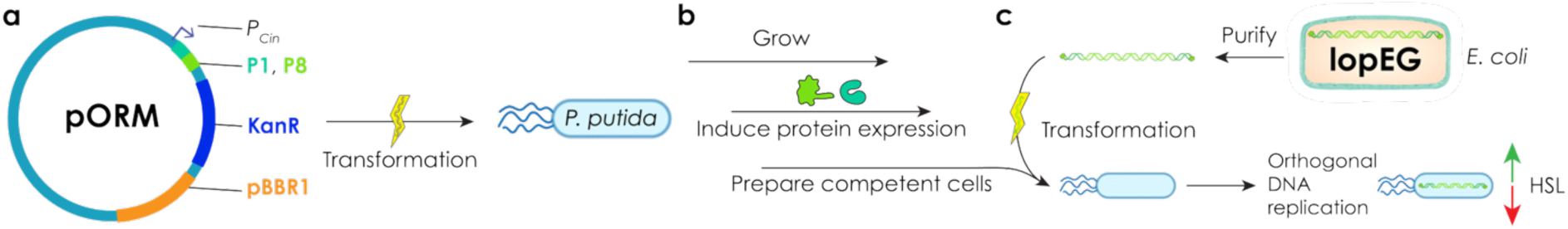
lopEG plasmids can be transformed into *P. putida*. **(a)** The antibiotic resistance on the pORM plasmid was changed to kanamycin to function in *P. putida*. **(b)** The *P. putida* culture is grown to the required OD, protein expression induced, and competent cells are prepared. **(c)** The lopEG plasmid is purified from *E. coli* cells and transformed in the freshly prepared *P. putida* competent cells. Plasmid copy number can subsequently be tuned using the OHC14-HSL inducer.

To this end, we modified pORM by replacing the chloramphenicol resistance gene with a kanamycin resistance marker, yielding pORM_KanR. This plasmid allowed us to circumvent the intrinsic chloramphenicol resistance of *P. putida*. *P. putida* electrocompetent cells were first transformed with pORM_KanR. Cells containing the plasmid where then made competent while inducing the expression of P1 and P8. lopEG plasmids from *E. coli* were subsequently transformed into these cells via electroporation. Using this workflow, we obtained transformants maintaining lopEG plasmids in *P. putida*. This result demonstrates that our platform is readily transferable to additional biotechnologically relevant species and highlight its potential for rapid gene evolution in alternative hosts.

### lopEG plasmid quantification in response to different HSL concentrations

To demonstrate that the lopEG plasmid copy number can be controlled, we grew *E. coli* and *P. putida* cells harbouring lopEG_GmR in liquid culture with different HSL concentrations. Notably, *P. putida* exhibited a higher sensitivity to the inducer, requiring only 1/10^th^ of the concentrations of HSL compared to *E. coli*. For both species, cells were harvested during stationary phase, and plasmid copy number was determined by qPCR using the single-copy chromosomal *dxs* gene as a reference for quantifying the lopEG-borne *GmR* marker (**Fig. 3** **and Supplementary Data 1**). In *E. coli*, the lopEG copy number increased approximately 67.4-fold, from ∼1.5 to ∼102.4 copies per cell (**Fig. 3a**)., whereas in *P. putida* it increased approximately 8.7-fold, from ∼7.7 to ∼67.3 copies per cell (**Fig. 3b**). Together, these results demonstrate that the orthogonal replicon copy number can be controlled over a wide range in both types of bacteria.

**Fig. 3:**
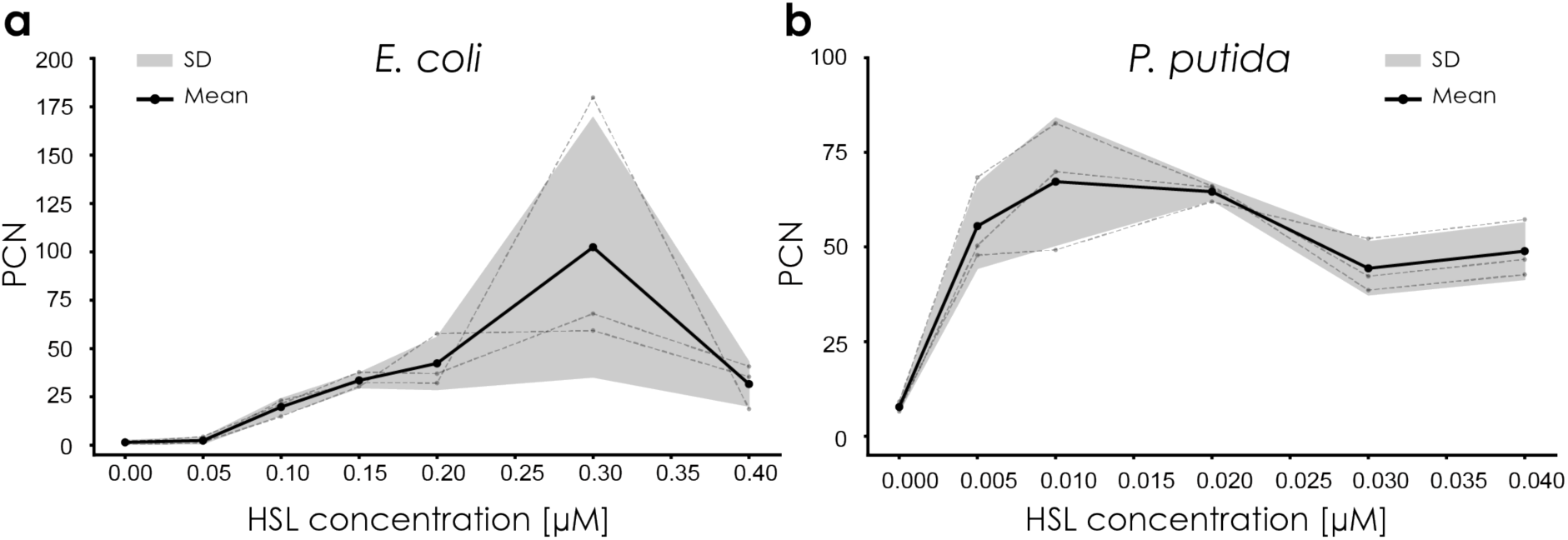
Control of lopEG plasmid copy number in *E. coli* and *P. putida*. **(a)** In *E. coli*, the lopEG plasmid copy number can be modulated over a broad dynamic range, from ∼1.5 to ∼102.4 copies per cell. **(b)** In *P. putida*, the lopEG plasmid copy number can be tuned from ∼7.7 to ∼67.3 copies per cell. In both panels, data are presented as mean ± standard deviation (SD). The black solid line and filled circles represent the mean of three biological replicates (n=3), individually shown as light grey dashed lines and circles. The grey shaded area surrounding the mean line indicates ± 1 SD. PCN = plasmid copy number.

### Orthogonal plasmid maintenance under different selective pressures

We next evaluated the stability of orthogonal plasmids in *E. coli* and *P. putida* over time (**Fig. 4**). To this end, cultures carrying the lopEG_GmR plasmid were grown under different selective conditions: (1) in the presence of gentamicin to maintain selection for lopEG, (2) in the presence of chloramphenicol or kanamycin to select for pORM or pORM_KanR respectively, or (3) in the absence of antibiotic selection. At each doubling time, cultures were plated on gentamicin-containing agar plates and the fraction of cells retaining the lopEG plasmid was determined relative to the initial population. In *E. coli*, when grown without selective pressure, more than 67% of cells lost the lopEG plasmid after only two generations, with only 3% of the population retaining the plasmid after 8 generations. When chloramphenicol was included to maintain the pORM plasmid, retention remained over 50% for more than 3 generations, with 27% of cells maintaining the plasmid over 8 generations. Under gentamicin selection, the plasmid was stably maintained over the course of the experiment (**Fig. 4a**). In *P. putida*, plasmid stability decreased in the absence of selective pressure, with 50% of cells losing the plasmid after two generations. Nevertheless, 18% of cells still retained lopEG after eight generations. In the presence of kanamycin, plasmid retention increased to 34%, after 8 generations, indicating slightly improved plasmid maintenance compared to *E. coli.* Under gentamicin selection, the lopEG plasmid was stably maintained across all generations, although some fluctuation between biological replicates was observed (**Fig. 4b**). Plate images for *E. coli* can be found in **Supplementary Fig. 1-3** whereas raw colony counts can be found in **Supplementary Table 1**. For *P. putida* Plate images can be found in **Supplementary Fig. 4-6** whereas raw colony counts can be found in **Supplementary Table 2**.

**Fig. 4:**
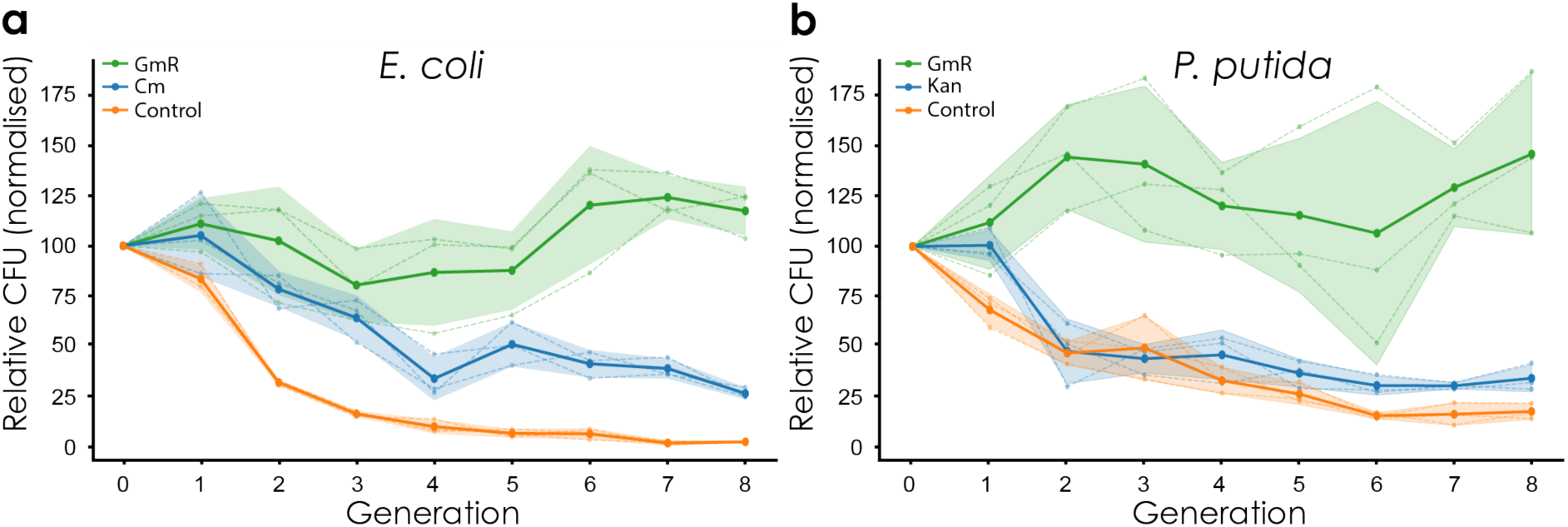
**Orthogonal plasmid maintenance under different selective pressures in *E. coli* and *P. putida*.**(**a**) Retention of the lopEG_GmR plasmid in *E. coli* during serial growth under gentamicin (GmR), chloramphenicol (Cm), or non-selective control conditions. After 8 generations, ∼27% and ∼3% plasmid retention was observed under Cm and control conditions, respectively, while plasmid maintenance remained stable under GmR selection. (**b**) Retention of the lopEG_GmR plasmid in *P. putida* during serial growth under gentamicin (GmR), kanamycin (Kan), or non-selective control conditions. After 8 generations, ∼34% and ∼18% plasmid retention was observed under Kan and control conditions, respectively, while stable plasmid maintenance was observed under GmR selection. In both panels, solid lines and large filled circles represent the mean of three biological replicates (n = 3). The individual replicates are shown as dashed lines with small circles, and the shaded band indicates ± 1 SD. CFU counts were normalised within each replicate to generation 0 (= 100%). CFU = colony-forming units.

### Evolution of an msfGFP expression construct yields an 11.8-fold increase in cell-normalised fluorescence

Finally, we tested whether PRIME could support the evolution of a simple target gene using only standard laboratory equipment and workflows. To this end, a variant of the lopEG plasmid carrying a gentamicin resistance gene and an in-frame *msfGFP*^21^ gene under the control of the BG17^21^ promoter was transformed in *E. coli* containing a pORM plasmid encoding an error-prone P1 polymerase variant (Y127A), which mutation rate was previously reported to be 5.61 × 10^−7^ s.p.b.^6^. Following transformation, the 10 brightest colonies were visually selected using a blue-light transilluminator, resuspended in liquid medium, and replated on large agar plates. This selection process was repeated seven times before sequencing the 30 brightest colonies obtained at the end of the experiment.

Sequencing revealed two independently evolved lopEG variants carrying mutations within the upstream regulatory regions (promoter and 5’ untranslated region) rather than the msfGFP coding sequence itself. Two further mutations were located within the in-frame gentamicin resistance gene (T9A and a silent mutation; **Supplementary Fig. 7**). The two selected lopEG variants were then retransformed into *E. coli* cells carrying a pORM plasmid encoding the wild-type DNA polymerase and single colonies were analysed in liquid culture using a microtitre plate reader. One evolved variant displayed a 1.5-fold increase in cell-normalised fluorescence relative to the starting construct, whereas the second variant exhibited an 11.8-fold increase (**Fig. 5 and supplementary Data 2**). Together, these results demonstrate that PRIME enables rapid *in vivo* evolution and selection of improved genetic constructs using simple and accessible experimental procedures.

**Figure 5:**
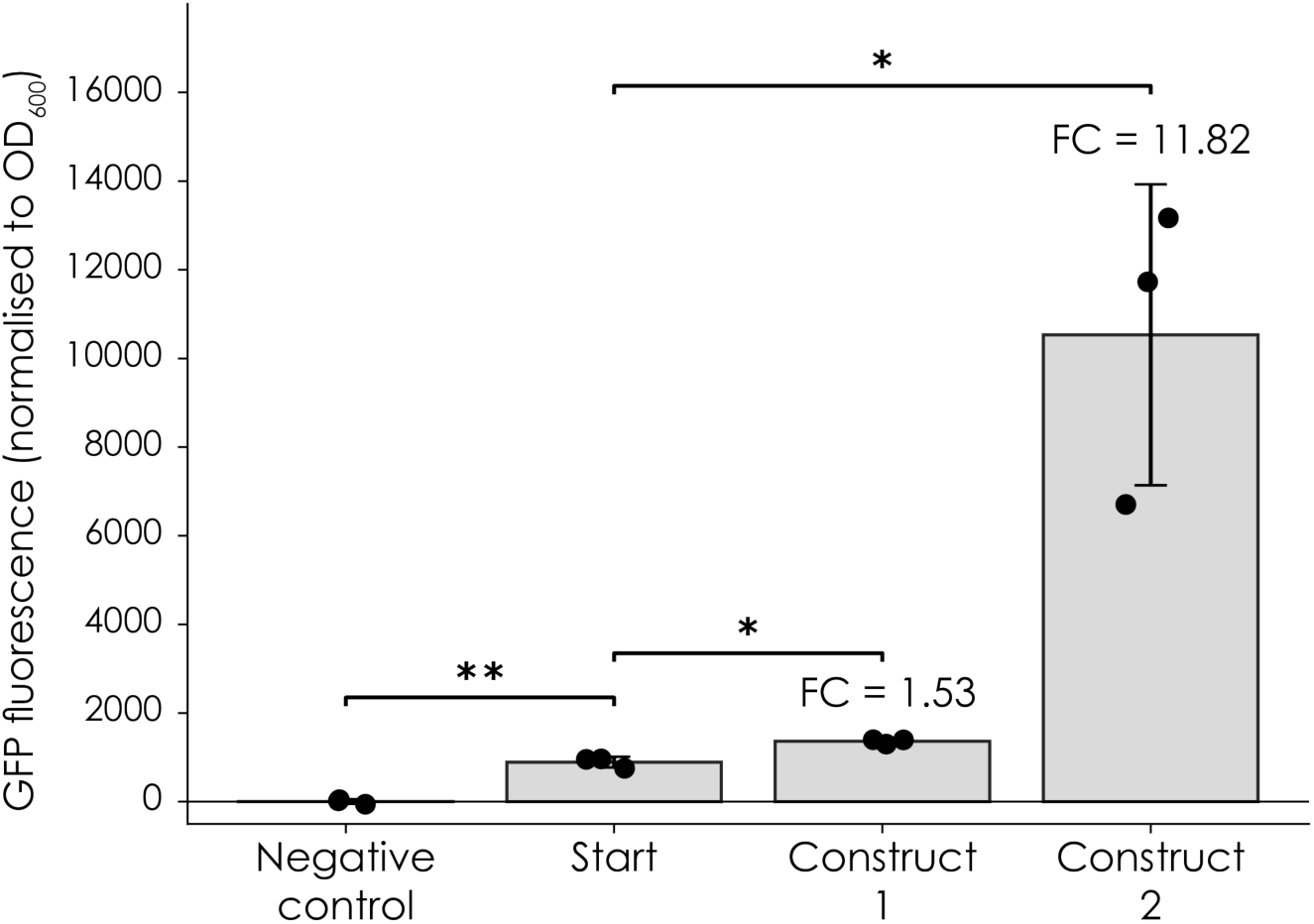
C**e**ll**-normalised fluorescence of evolved constructs relative to the starting construct.** Background-corrected GFP fluorescence was normalised to cell density using background-corrected OD_600_ measurements. For each biological replicate, three measurements with background-corrected OD_600_ values closest to 0.075 were selected: The measurement closest to OD_600_ = 0.075 and the nearest measurements above and below this value. An OD_600_ of 0.075 was selected as it corresponds to the exponential growth phase, providing a consistent and biologically comparable point for assessing fluorescence across replicates. For each of these measurements, GFP fluorescence was divided by the corresponding background-corrected OD_600_ to obtain cell-normalised fluorescence. The mean of these three values was calculated for each biological replicate and data are shown as mean ± SD from three biological replicates (n = 3). Individual biological replicate values are indicated by black dots. The negative control consisted of a replicon carrying the gentamicin resistance marker but lacking the *msfGFP* reporter gene. Fold-change (FC) values are shown relative to the starting construct (Start). Statistical comparisons of normalised fluorescence were performed against the starting construct using an unpaired two-tailed Welch’s *t*-test, which does not assume equal variances. No adjustment for multiple comparisons was applied, with horizontal brackets indicating pairwise comparisons. The negative control showed significantly lower normalised fluorescence relative to the starting construct (*t*(2.77) = −11.72, *p* = 0.0019). Evolved construct 1 exhibited a 1.5-fold increase in normalised fluorescence relative to the starting construct (*t*(2.83) = 6.19, *p* = 0.0100), whereas evolved construct 2 exhibited an 11.8-fold increase (*t*(2.00) = 4.92, *p* = 0.0387). Significance levels are indicated as follows: *p* < 0.05 (*), *p* < 0.01 (**), and *p* < 0.001 (***). Raw data can be found in **Supplementary Data 2**.

## Discussion

The work presented here establishes PRIME (<u>P</u>rotein-primed <u>R</u>eplication for *In vivo* <u>M</u>utagenesis and <u>E</u>volution), a workflow for the facile implementation of orthogonal plasmids replication in two different biotechnologically relevant organisms, *E. coli* and *P. putida*. Given the broad host-range of the PRD1 bacteriophage, we anticipate that this approach could be extended to further organisms. A key advantage of PRIME is that it does not require genomic modification, relying instead on a broad-host-range plasmid system and accessory plasmids for transformation and replication initiation. This greatly simplifies the deployment of orthogonal replicons in established laboratory strains, metabolically engineered production hosts, and bacterial chassis where genome engineering is inefficient or poorly standardised.

Using an error-prone variant of the P1 polymerase^6^, we demonstrate that constructs encoded on the lopEG orthogonal plasmid can be subjected to rapid continuous evolution *in vivo*. As a proof of concept, we targeted a construct containing an msfGFP reporter for enhanced fluorescence, achieving an 11.8-fold increase after 7 iterative cycles of diversification and visual colony-based screening. These results establish PRIME as a streamlined platform for continuous directed evolution using minimal infrastructure and standard microbiological workflows.

Furthermore, PRIME enables the control of orthogonal plasmid copy number in both *E. coli* and *P. putida* through inducible expression of the PRD1 replication machinery. This provides a modular framework in which replication dynamics can be matched to host physiology, target gene burden and selection strategy.

Our data further clarify the role of the PRD1 single-stranded DNA-binding protein P12^9,22^. We had previously shown that P12 strongly binds to ssDNA and protects it against nuclease activity^9^. In contrary to P19, which had been shown to be dispensable for PRD1 bacteriophage propagation^12^ but helpful for DNA transformation^6^, P12 was shown to be required for efficient transformation^6^. Our results confirm that both single-stranded DNA binding protein (SSBs) supports efficient transformation of linear DNA but further reveal that P12 is also dispensable for maintenance of established orthogonal plasmids. We propose that P12 protects exposed DNA immediately following transformation, prior to terminal protein P8-mediated capping. Once protein-primed DNA replication is established, P12 is no longer required and lopEG plasmids can be stably maintained by P1 and P8 alone. These findings suggest partial functional redundancy with host single-stranded DNA-binding proteins, such as *E. coli* SSB^23^ during replication but not during DNA uptake.

Importantly, the removal of P12 from the final pORM plasmid allows PRIME to circumvent the toxic effects of P12 reported in the literature^10–12^. We previously showed that this toxicity is linked to the ssDNA-binding activity of P12^9^. Although P12 has been shown to be crucial for PRD1 bacteriophage propagation^12^, our observation that it is dispensable for DNA replication suggests that its critical function during phage replication likely extends beyond direct support of DNA synthesis. Consequently, these findings open new avenues of inquiry into how P12 functions within the broader phage replication cycle.

Collectively, this work establishes a portable, plasmid-based orthogonal replication platform that expands access to continuous directed evolution technologies beyond specialist laboratories. Because the system relies on standard cloning and cultivation procedures, it is readily deployable across diverse experimental settings. In particular, its compatibility with metabolically engineered strains makes it well suited for the optimisation of biosynthetic pathways through iterative *in vivo* evolution of individual enzymatic steps. We therefore envision that PRIME will facilitate the implementation of continuous directed evolution approaches across a broad range of laboratories and experimental settings.

## Methods

### Cloning of the plasmid containing the Orthogonal Replication Machinery (pORM)

The pORM plasmid was constructed stepwise. First, the replication origin of pAJM1642 (Addgene #108535) was replaced with the pBBR1 origin. The vector backbone was amplified using primers pAJM1642_SapI_fw and pAJM1642_SapI_rev, and the pBBR1 origin using primers pBBR1_SapI_fw and pBBR1_SapI_rev. PCR products were digested with SapI restriction enzyme (New England Biolabs), ligated (Roche), and transformed into DH10B cells. The antibiotic resistance cassette was subsequently switched from kanamycin to chloramphenicol by amplifying the plasmid backbone with primers pAJM1642-KanR_NotI_fw and pAJM1642-KanR_XbaI_rev, and the chloramphenicol acetyltransferase gene with primers CAT_XbaI_fw and CAT_NotI_rev. PCR products were digested with XbaI and NotI restriction enzymes (New England Biolabs), ligated, and transformed into DH10B cells, yielding pORM_CAT_YFP. The YFP gene was then replaced with the operon encoding for the orthogonal replication machinery by digestion with BsaI-HF-v2 and KpnI-HF restriction enzymes (New England Biolabs) and ligation with a similarly digested insert, yielding pORM_P12_P19. P12 variants were introduced in this vector by amplifying the plasmid backbone using primers pORM-P12_SapI_fw and pORM-P12_SapI_rev. The P12_T2 variant was amplified using primer P12T2_SapI_fw and P12T2 _SapI_rev. PCR products were digested with SapI restriction enzyme, ligated, and transformed into DH10B cells, yielding pORM_P12_T2. The plasmid pORM was generated by removing the *p12* and *p19* genes using primers P12&P19_remove_SapI_fw and P12&P19_remove_SapI_rev, followed by SapI digestion and transformation into DH10B cells. To construct pORM_KanR, the kanamycin resistance gene was amplified using primers KanR_for_pORM_XbaI_fw and KanR_for_pORM_BsrGI_rev. Both the PCR product and the pORM backbone were digested with XbaI and BsrGI restriction enzymes (New England Biolabs), followed by ligation and transformation into DH10B cells. Finally, the Y127A mutation was introduced onto P1 to generate an error-prone polymerase using primers pORM_P1Y127A_SapI_fw and pORM_P1Y127A_SapI_rev, yielding pORM_P1Y127A. Primer sequences can be found in **Supplementary Table 3** whereas plasmid maps are present in **Supplementary Data 3**.

### Cloning of the Orthogonal Transformation Helper plasmid (pOTH)

The pOTH plasmid was assembled from two PCR fragments. The first fragment, containing the pSC101(ts) replication origin, the L-arabinose operon, and the *gam* gene under the control of the *p_BAD_* promoter, was amplified using primers pOTH_bb_BsaI_fw and pOTH_bb_BsaI_rev. Codon-optimised sequences encoding P12 and P19 (GeneArt, Thermo Fisher Scientific) were amplified using primers p12CO_pOTH_BsaI_fw and p19CO_pOTH_BsaI_rev. Both fragments were digested with BsaI-HF-v2 restriction enzyme, ligated (Roche), and transformed into DH10B cells, yielding pOTH. Primer sequences can be found in **Supplementary Table 3** whereas plasmid maps are present in **Supplementary Data 3**.

### Cloning of the plasmid for the production Orthogonal Replicons Flanked by Inverted Terminal Repeats (pORFITR)

A plasmid containing a synthetic gene construct consisting of a gentamicin resistance gene and *msfGFP* under control of the BG17^21^ promoter and flanked by ITRs sequences (ordered at GENEWIZ, Azenta) was digested with XhoI and NheI restriction enzymes (New England Biolabs) and gel-purified using a NucleoSpin Gel and PCR Clean-up kit (Macherey-Nagel). In parallel, a pEMG plasmid^20^ was digested with the same restriction enzymes and dephosphorylated (Roche). The fragments were ligated and transformed into DH5αλpir cells^20^, yielding pORFITR_BG17_GmR_msfGFP. Promoter variants were generated using primers pORFITR_PromChange_AvrII_fw, and pORFITR_BG17_AvrII_rev or pORFITR_BG28_AvrII_rev. The PCR product was digested with AvrII restriction enzyme (New England Biolabs), ligated and transformed in DH10B cells.

To construct pORFITR_BG17_GmR, the *GmR* gene was amplified from the pORFITR_BG17_GmR_msfGFP plasmid using primers GmR_Bpu10I_fw and GmR_SbfI_rev. The PCR product and pORFITR backbone were digested with Bpu10I and SbfI restriction enzymes (New England Biolabs), followed by dephosphorylation of the vector backbone. Ligation and transformation into DH5αλpir cells, yielded pORFITR_BG17_GmR. Primer sequences can be found in **Supplementary Table 3** whereas plasmid maps are present in **Supplementary Data 3**.

### Production and purification of Linear Orthogonal Plasmid with Evolving Genes (lopEG)

The lopEG plasmids were generated by digestion of the different pORFITR plasmids with MlyI restriction enzyme (New England Biolabs). The resulting digested DNA was either dephosphorylated or left untreated, and subsequently either resolved on a 1% agarose gel followed by band extraction using the NucleoSpin Gel and PCR Clean-up kit (Macherey-Nagel), or directly purified using the same kit without prior gel separation, before transformation.

lopEG plasmids were purified from *E. coli* cells by using the standard protocol of the NucleoSpin Plasmid Mini kit for plasmid DNA (Macherey-Nagel).

### Transformation of lopEG plasmids in *E. coli*

DH10B cells were first transformed with the pOTH plasmid and plated overnight at 30 °C. Electrocompetent cells were prepared from a single colony by four successive washes with cold deionised water. These cells were then transformed with the pORM plasmid and plated overnight at 30 °C. A colony of doubly transformed cells was then picked and grown overnight in LB medium supplemented with chloramphenicol (34 µg/mL) and tetracycline (3 µg/mL) at 30 °C with constant shaking. The following morning, this culture was used to inoculate fresh LB medium (1:100) containing the same antibiotics. Cells were grown at 30 °C with shaking until an optical density at 600 nm (OD_600_) of 0.3 was reached, at which point L-arabinose (0.4%) and 0.2 µM OHC14-HSL inducer (HSL)^13^ were added to induce protein expression. Cultures moved to 37 °C and incubated for 45 min, chilled on ice for 15 min, and harvested by centrifugation (4000 × *g*, 5 min, 4 °C). Cell pellets were washed four times with pre-chilled deionised water to generate electrocompetent cells, which were then transformed with 1 µg of predigested pORFITR plasmids using pre-chilled 2 mm electroporation cuvettes (Cell Projects Ltd). Electroporation was performed with a single 2500 V pulse using a MicroPulser Electroporator (Bio-Rad). Cells were resuspended in SOC medium supplemented with 0.1 µM HSL and incubated at 37 °C with shaking at 800 rpm for recovery prior to plating LB-agar plates supplemented with gentamicin (20 µg/mL; Sigma-Aldrich).

### Transformation of lopEG plasmids in *P. putida*

Electrocompetent *P. putida* KT2440 cells^24^ were prepared from an overnight culture in LB medium by three successive washes with pre-chilled 300 mM sucrose, each followed by centrifugation. Cells were electroporated with pORM_KanR using pre-chilled 2 mm cuvettes (Cell Projects Ltd) and a single pulse at 2500 V (MicroPulser, Bio-Rad). Following electroporation, cells were resuspended in SOC medium and incubated at 30 °C with shaking (800 rpm) for recovery prior to plating on cetrimide agar supplemented with 50 µg/mL kanamycin. A colony was then picked and grown overnight at 30 °C with constant shaking in LB medium supplemented with kanamycin (50 µg/mL). The following morning, this culture was used to inoculate fresh LB medium (1:100) containing the same antibiotic. Cells were grown at 30 °C with shaking until an optical density at 600 nm (OD_600_) of 0.5 was reached, at which point 0.05 µM HSL was added to induce protein expression. The culture was incubated for 1h at 30 °C before being chilled on ice for 15 min, and harvested by centrifugation (4000 × *g*, 5 min, 4 °C). Cell pellets were washed three times with a pre-chilled 300 mM sucrose solution to generate electrocompetent cells. Cells were then transformed with lopEG_GmR plasmid purified from *E. coli* cultures grown in the presence of 0.3 µM HSL using a NucleoSpin Plasmid Mini kit for plasmid DNA (Macherey-Nagel). The plasmid was transformed using a single 2500 V pulse and cells allowed to recover in SOC medium supplemented with 0.01 µM HSL at 30 °C with shaking (800 rpm) for one hour prior to plating on cetrimide agar supplemented with gentamicin (20 µg/mL).

### Orthogonal plasmid maintenance quantification

For *E. coli*, DH10B cells were first transformed with the pOTH and pORM plasmids, followed by transformation with the lopEG_GmR plasmid. Cells were plated on LB agar supplemented with gentamicin (20 µg/mL) and 0.1 µM HSL. Three independent colonies were inoculated into LB medium containing gentamicin (20 µg/mL) and 0.1 µM HSL and grown overnight at 37 °C with shaking (220 r.p.m.). The following day, cultures were washed and diluted three times into fresh medium containing either (1) gentamicin (10 µg/mL) and 0.1 µM HSL, (2) chloramphenicol (34 µg/mL) and 0.1 µM HSL, or (3) 0.1 µM HSL only. Cultures were adjusted to OD_600_ = 0.25 and grown at 37 °C with shaking (220 r.p.m.) to OD_600_ = 0.5. Cultures were then diluted back to OD_600_ = 0.25 and regrown to OD_600_ = 0.5. This serial propagation procedure was repeated for a total of eight doubling time. At the initial time point and after each doubling, approximately 500 cells from each culture were plated on LB agar supplemented with gentamicin (10 µg/mL) and 0.1 µM HSL. Images of the plates were captured using a digital camera and colonies were counted using the Colony Counter Plugin for ImageJ v. 1.54f^25^ authored by Bruno Vieira, in order to determine the fraction of cells retaining the lopEG_GmR plasmid over time (images of the plates can be found in **Supplementary Fig. 1-3)**.

For *P. putida*, cells carrying pORM_KanR were transformed with lopEG_GmR plasmid DNA purified from *E. coli*. Transformants were plated on LB agar supplemented with gentamicin (10 µg/mL) and 0.01 µM HSL. Three independent colonies were inoculated into LB medium containing gentamicin (20 µg/mL) and 0.01 µM HSL and grown overnight at 30 °C with shaking (220 r.p.m.). The following day, cultures were washed and diluted into fresh medium containing either (1) gentamicin (10 µg/mL) and 0.01 µM HSL, (2) kanamycin (50 µg/mL) and 0.01 µM HSL, or (3) 0.01 µM HSL only. Cultures were adjusted to OD_600_ = 0.25 and grown at 30 °C with shaking (220 r.p.m.) to OD_600_ = 0.5. Cultures were subsequently diluted back to OD_600_ = 0.25 and regrown to OD_600_ = 0.5. This propagation cycle was repeated for a total of eight doublings. At the initial time point and after each doubling, approximately 500 cells from each culture were plated on LB agar supplemented with gentamicin (10 µg/mL) and 0.01 µM HSL. Images of the plates were captured using a digital camera and colonies were counted using the Colony Counter Plugin for ImageJ v. 1.54f^25^ authored by Bruno Vieira, in order to determine the fraction of cells retaining the lopEG_GmR plasmid over time (images of the plates can be found in **Supplementary Fig. 4-6)**.

### Determination of plasmid copy number by qPCR

Aliquots containing ∼10⁸ cells, and ∼2 × 10⁸ cells for *P. putida*, were pelleted by centrifugation at 1,700 × g for 3 min and washed once with 0.5 mL of sterile water. Pellets were resuspended in QuickExtract Bacterial DNA Extraction Solution (Lucigen) and incubated at room temperature for 15 min. Quantitative PCR was performed using Fast SYBR Green 2X Master Mix (Applied Biosystems) on a Roche LightCycler 96. Three independent biological replicates were analysed, each measured in technical triplicates. Reactions were prepared in a final volume of 20 µL with 1 µL of 1:10 diluted cell lysate. Primers targeting the single-copy chromosomal *dxs* gene and the plasmid-borne *GmR* gene were used (sequences are listed in **Supplementary Table 3**). Primer efficiencies were evaluated using a dilution series of *E. coli* DH10B and *P. putida* KT2440 for *dxs,* and purified plasmid DNA for *GmR* . The PCR cycle threshold (Ct) values were extracted from the raw fluorescence data using the LightCycler software. The plasmid copy number (PCN) was calculated as^26^ :

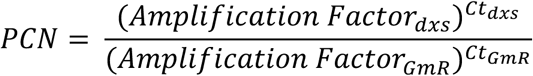

### msfGFP evolution

Cells were first transformed with a version of the pORM plasmid carrying the Y127A error-prone variant of P1^6^ and the pOTH plasmid. These cells were subsequently transformed with the lopEG_BG17_GmR_msfGFP plasmid and plated on LB-agar plates containing gentamicin (20 µg/mL) and 0.1 µM HSL. Ten fluorescent colonies were selected and pooled in liquid medium. From this solution, a subset of 5000 cells was then spread onto large LB agar plates containing gentamicin (10 µg/mL) and 0.2 µM HSL and incubated overnight at 37 °C. The next day, colony fluorescence was screened visually under blue-light illumination. Ten colonies displaying the highest fluorescence were selected and subjected to subsequent rounds of diversification and screening following the same procedure. This iterative evolution workflow was repeated for 7 rounds.

### msfGFP fluorescence measurements

lopEG plasmid containing evolved GmR_msfGFP cassettes were extracted from cells obtained after 7 rounds of evolution. Purified lopEG plasmids were transformed into *E. coli* DH10B cells carrying the pORM plasmid encoding the wild-type P1 polymerase and plated on LB agar supplemented with gentamicin (20 µg/mL) and 0.1 µM HSL. For each evolved construct, three independent colonies were inoculated into 100 µL LB medium supplemented with gentamicin (10 µg/mL) and 0.1 µM HSL in a black clear-bottom 96-well microplate (Nunc™, Thermo Fisher Scientific). Cultures were incubated at 37 °C in an Infinite M200 Pro plate reader (Tecan) under continuous shaking. GFP fluorescence (λ_ex_ = 485 nm; λ_em_ = 535 nm) and OD_600_ were monitored throughout growth. Fluorescence values were normalised to OD_600_ to determine fluorescence per cell.

## Statistics and Reproducibility

No statistical methods were used to predetermine sample size. Sample sizes were chosen in accordance with standard practice in the field for such experiments. All experiments were performed with n = 3 biological replicates. Biological replicates were defined as independent colonies picked from a single transformation. For the plasmid maintenance experiments (**Fig. 4**), each biological replicate corresponds to an independent lineage propagated in parallel over all nine sampling points (generations 0–8). Technical replicates, where used, refer to repeated measurements of the same culture. No data points or replicates were excluded from the analyses. Statistical analyses were performed in Python (v3.10.11) using NumPy (v2.1.3) and SciPy (v1.15.3). Pairwise comparisons of normalised GFP fluorescence (**Fig. 5**) were assessed using an unpaired two-tailed Welch’s *t*-test, assuming unequal variances, as implemented in scipy.stats.ttest_ind, without adjustment for multiple comparisons. Exact *p*-values are reported in the corresponding figure legends. Data are presented as mean ± SD unless otherwise stated, with all individual biological replicates displayed.

## Data availability

All data supporting the findings of this study are available within the article and its Supplementary Information files.

## Biological materials availability

Plasmids and bacterial strains generated in this study are available from the corresponding author upon request.

## Supporting information

Description of Additional Supplemental Files

## Acknowledgement

We thank S. Panke for helpful discussions and sharing of facilities. We are also grateful to P. Marlière for initial discussions on PRD1.

## Authors contributions

S.S. and N.H.-D. cloned the original set of PRIME plasmids. L.V., L.K.T and N.H.-D. cloned the improved set of PRIME plasmids. E.D, L.V. and N.H.-D. characterised the system. M.O. provided pEMG, *Pseudomonas putida* KT2440 and guidance on performing experiments related to this strain. N.H.-D. performed experiments with *P. putida*. L.K.T. performed the plasmid quantification experiments. N.H.-D. supervised E.D., S.S., L.V. and L.K.T. and defined the direction of research. All authors analysed and interpreted the data. N.H.-D. and L.K.T. wrote the manuscript with input from all authors.

## Additional information

### Supplementary information

The online version contains supplementary material available at xxx

**Correspondence** and requests for materials should be addressed to Nicolas Huguenin-Dezot.

