## Supplementary material for "PRIME: A modular plasmid-based platform for continuous in vivo evolution across bacteria": Description of Additional Supplemental Files

### The following files are provided separately from the present document:

**Supplementary Data 1**: Raw qPCR Ct values

**Supplementary Data 2**: Biomass and fluorescence measurements of selected BG17_GmR_msfGFP constructs

**Supplementary Data 3**: Plasmid maps of plasmids described in this study
