## Supplemental Information for "PRIME: A modular plasmid-based platform for continuous in vivo evolution across bacteria"

### Supplementary information

#### Table of content:


**Page 2:** **Supplementary Fig. 1**: Plate images of *Escherichia coli* grown in the presence of gentamicin.

**Page 3:** **Supplementary Fig. 2**: Plate images of *Escherichia coli* grown in the presence of chloramphenicol.

**Page 4:** **Supplementary Fig. 3**: Plate images of *Escherichia coli* grown in the absence of antibiotic.

**Page 5:** **Supplementary Fig. 4**: Plate images for *Pseudomonas putida* grown in the presence of gentamicin.

**Page 6:** **Supplementary Fig. 5**: Plate images for *Pseudomonas putida* grown in the presence of kanamycin.

**Page 7:** **Supplementary Fig. 6**: Plate images for *Pseudomonas putida* grown in the absence of antibiotic.

**Page 8-9:** **Supplementary Fig. 7**: Mutations found in lopEG_BG17_GmR_msfGFP plasmids after continuous directed evolution.

**Page 10:** **Supplementary Table 1**: Raw colony counts for *Escherichia coli*.

**Page 11:** **Supplementary Table 2**: Raw colony counts for *Pseudomonas putida*.

**Page 12:** **Supplementary Table 3**: Primers used in this study.


#### The following files are provided separately from the present document:

**Supplementary Data 1**: Raw qPCR Ct values

**Supplementary Data 2**: Biomass and fluorescence measurements of selected BG17_GmR_msfGFP constructs

**Supplementary Data 3**: Plasmid maps of plasmids described in this study

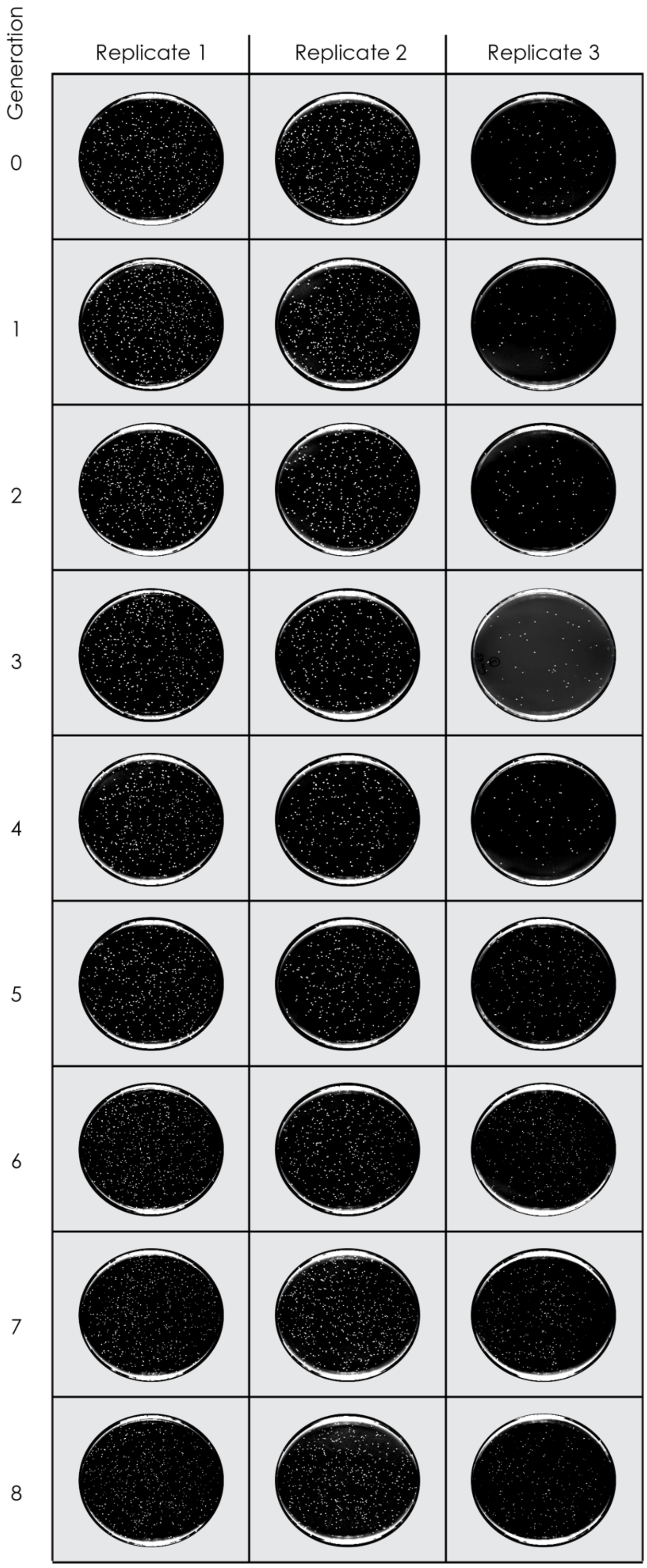


**Supplementary Fig. 1**: Plate images of *Escherichia coli* grown in the presence of gentamicin. Images were acquired under high-contrast illumination conditions. Three biological replicates were performed for each generation.

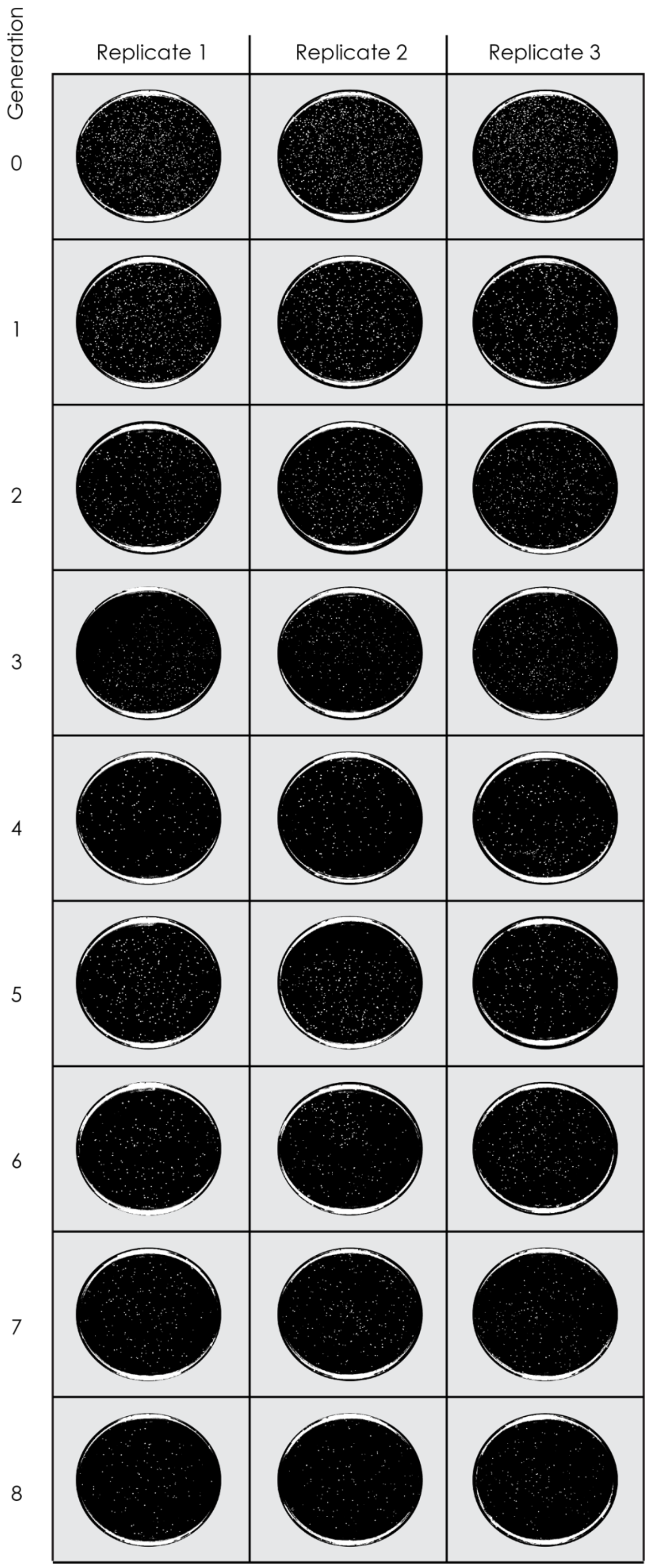


**Supplementary Fig. 2**: Plate images of *Escherichia coli* grown in the presence of chloramphenicol. Images were acquired under high-contrast illumination conditions. Three biological replicates were performed for each generation. The generation 0 sample inadvertently contained twice the intended number of cells. This was taken into account in subsequent calculations.

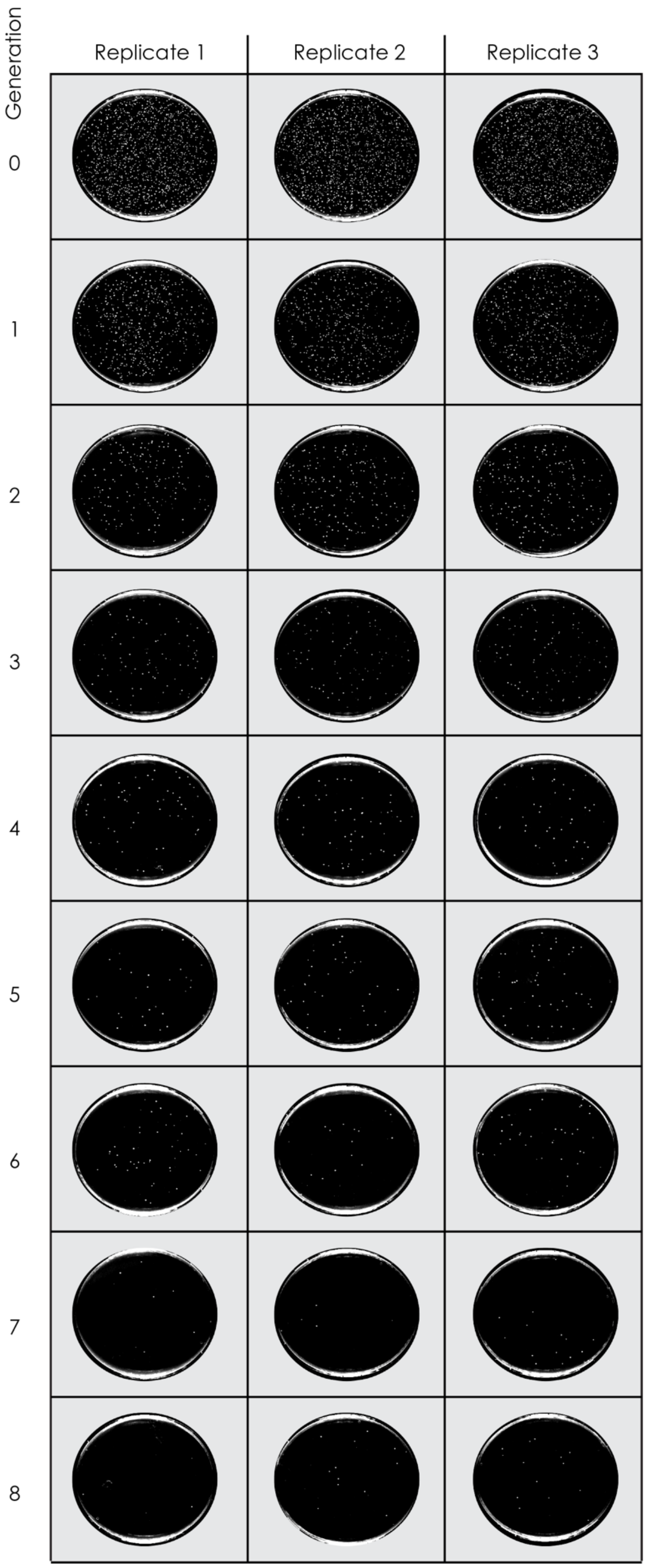


**Supplementary Fig. 3**: Plate images of *Escherichia coli* grown in the absence of antibiotic. Images were acquired under high-contrast illumination conditions. Three biological replicates were performed for each generation. The generation 0 sample inadvertently contained twice the intended number of cells. This was taken into account in subsequent calculations.

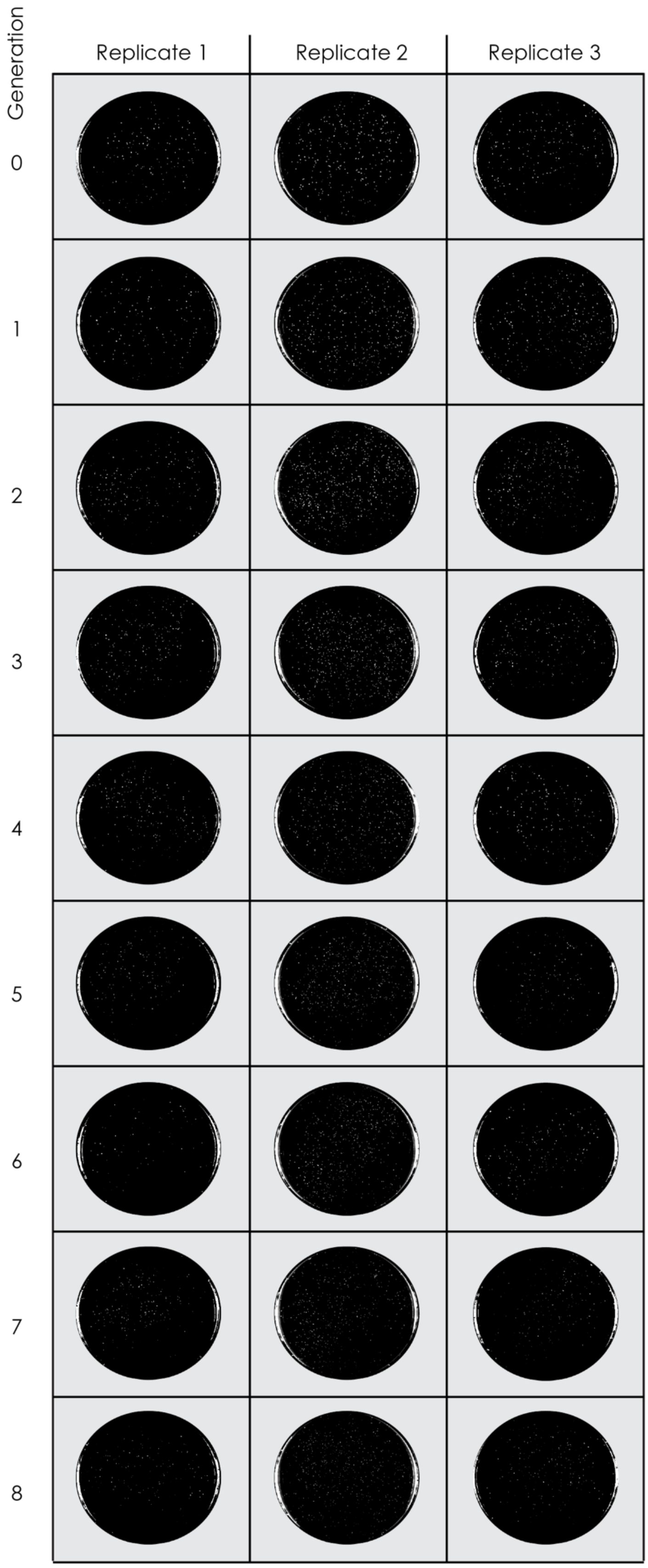


**Supplementary Fig. 4**: Plate images for *Pseudomonas putida* grown in the presence of gentamicin. Images were acquired under high-contrast illumination conditions. Three biological replicates were performed for each generation.

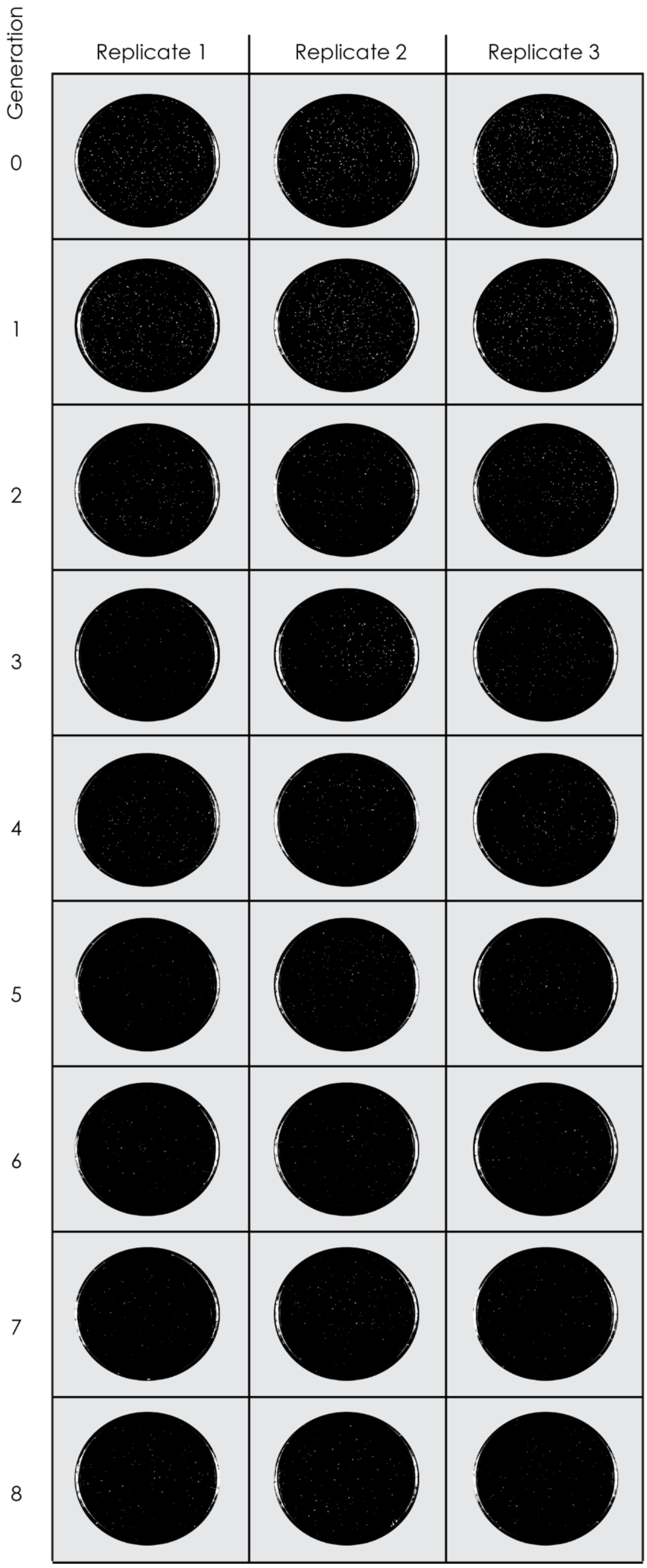


**Supplementary Fig. 5**: Plate images for *Pseudomonas putida* grown in the presence of kanamycin. Images were acquired under high-contrast illumination conditions. Three biological replicates were performed for each generation.

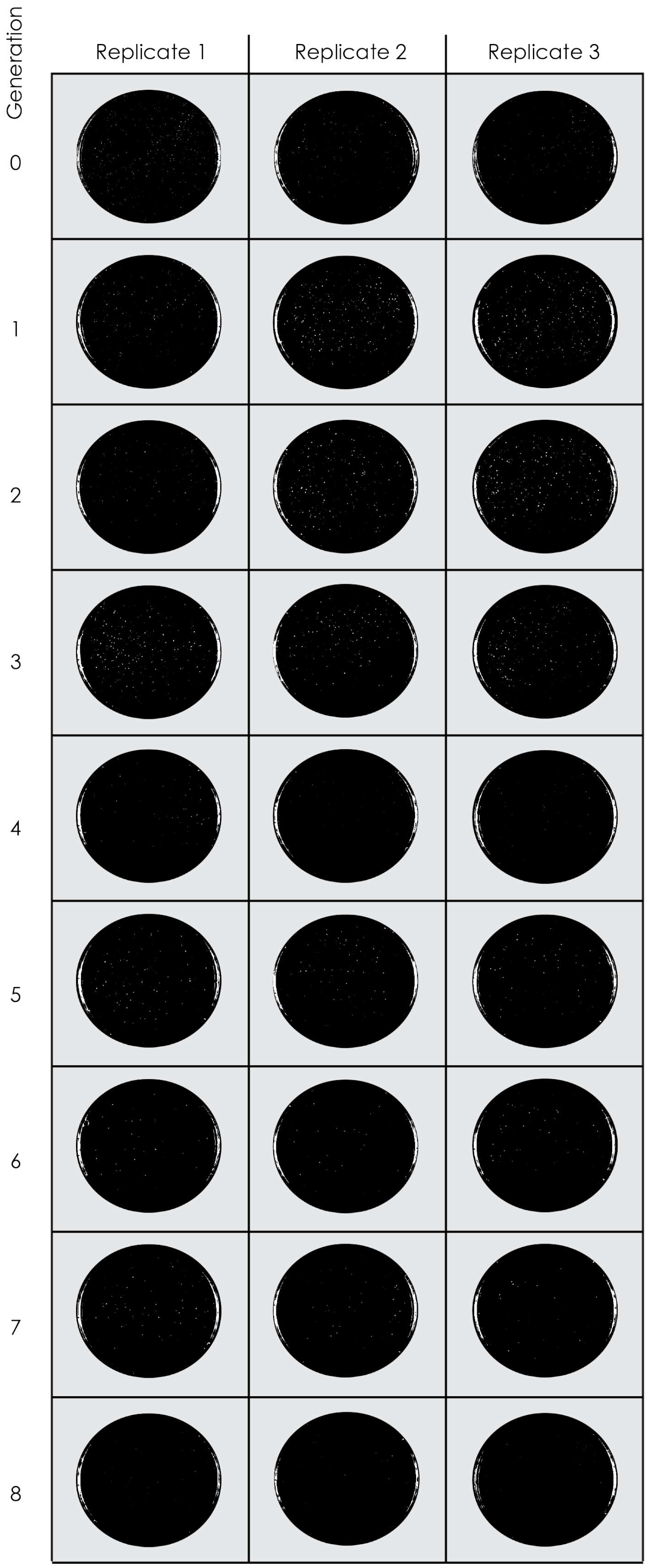


**Supplementary Fig. 6**: Plate images for *Pseudomonas putida* grown in the absence of antibiotic. Images were acquired under high-contrast illumination conditions. Three biological replicates were performed for each generation.

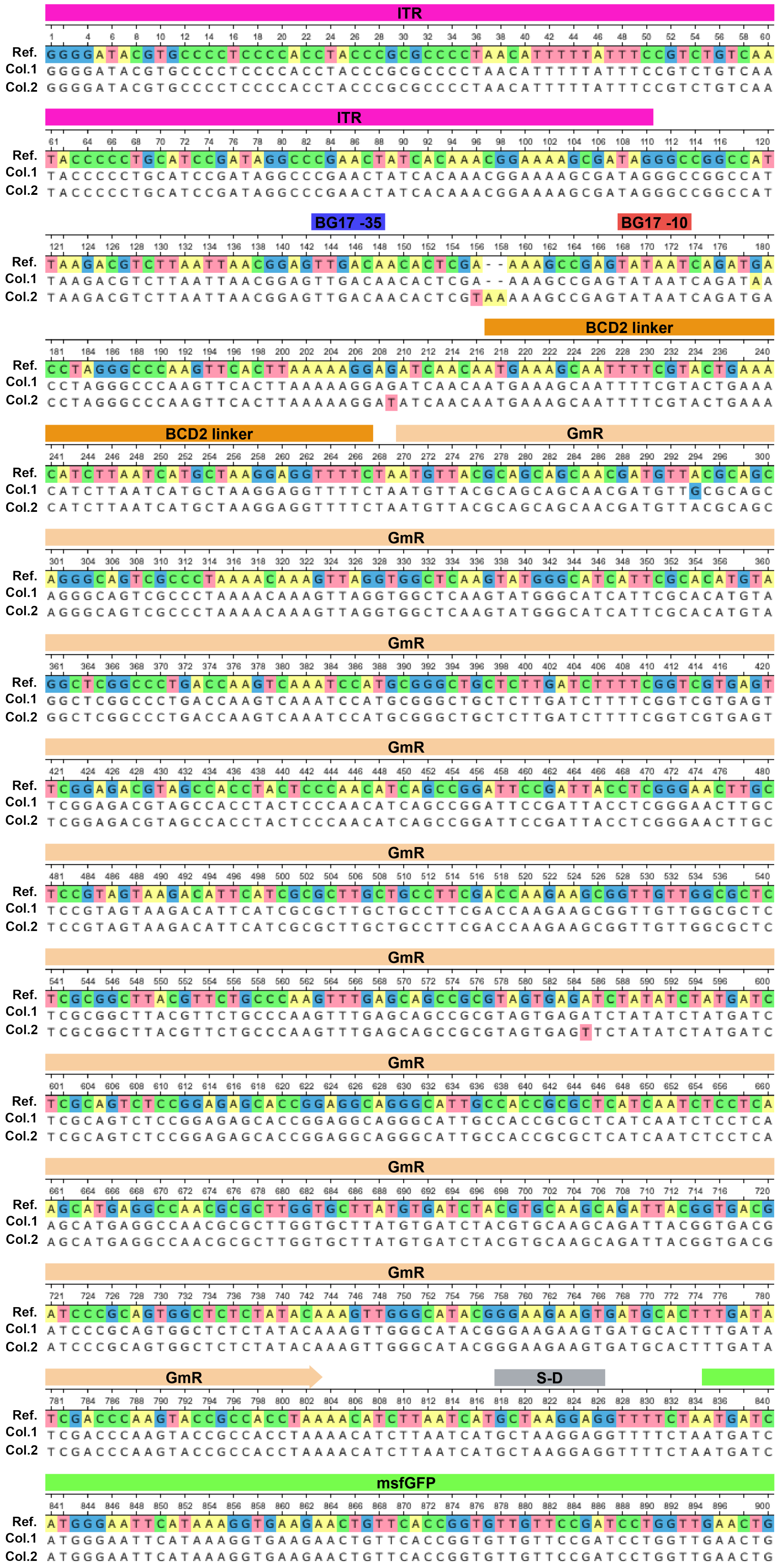
ITR
BG17 -35
BG17 -10
BCD2 linker
GmR
S-D
msfGFP
Ref.
Col.1
Col.2

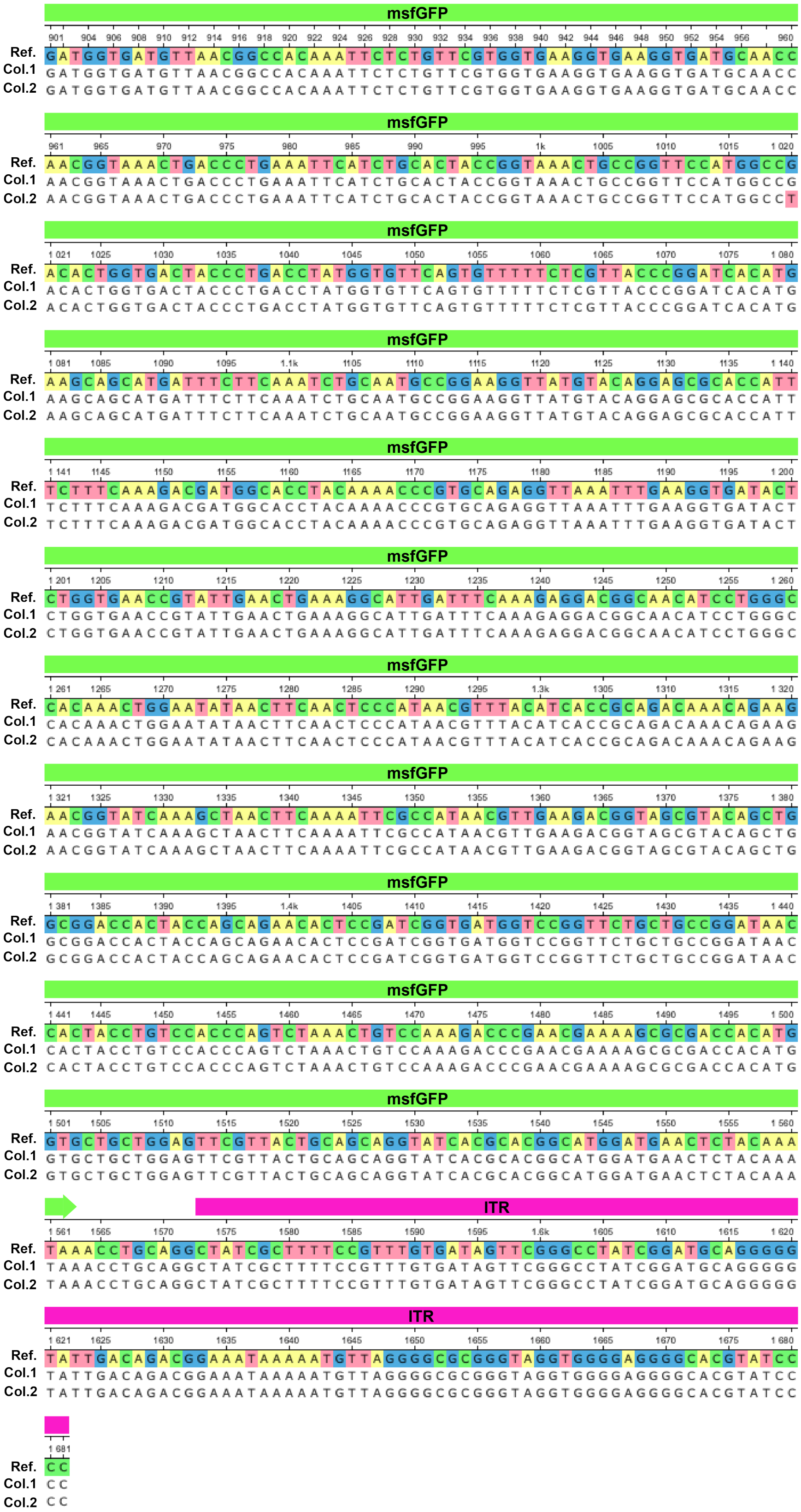

**Supplementary Fig. 7**: Mutations found in lopEG_BG17_GmR_msfGFP plasmids after continuous directed evolution. Features are colour-coded at the top of the sequence and mutation highlighted.

**Supplementary Table 1**: Raw colony counts for *Escherichia coli*. Raw colony counts were acquired from plate pictures using Colony Counter plugin implemented in Fiji v1.54f.

| *E. coli* | **Gentamicin** | | | **Chloramphenicol** | | | **No selection pressure** | | |
|---|---|---|---|---|---|---|---|---|---|
| **Generation** | **1** | **2** | **3** | **1** | **2** | **3** | **1** | **2** | **3** |
| **0** | 391 | 439 | 301 | 517 | 470 | 524 | 598 | 579 | 597 |
| **1** | 473 | 426 | 346 | 654 | 483 | 452 | 478 | 527 | 477 |
| **2** | 461 | 315 | 355 | 356 | 383 | 447 | 196 | 178 | 197 |
| **3** | 386 | 277 | 240 | 377 | 319 | 273 | 100 | 88 | 106 |
| **4** | 404 | 248 | 303 | 239 | 127 | 152 | 47 | 81 | 53 |
| **5** | 385 | 288 | 299 | 261 | 292 | 213 | 54 | 29 | 40 |
| **6** | 539 | 380 | 410 | 177 | 202 | 247 | 43 | 52 | 22 |
| **7** | 533 | 521 | 353 | 188 | 210 | 190 | 19 | 6 | 12 |
| **8** | 484 | 455 | 375 | 130 | 118 | 156 | 13 | 17 | 17 |

**Supplementary Table 2**: Raw colony counts for *Pseudomonas putida*. Raw colony counts were acquired from plate pictures using Colony Counter plugin implemented in Fiji v1.54f.

| *P. putida* | **Gentamicin** | | | **Kanamycin** | | | **No selection pressure** | | |
|---|---|---|---|---|---|---|---|---|---|
| **Generation** | **1** | **2** | **3** | **1** | **2** | **3** | **1** | **2** | **3** |
| **0** | 216 | 324 | 258 | 359 | 508 | 348 | 585 | 722 | 637 |
| **1** | 185 | 390 | 335 | 347 | 487 | 380 | 434 | 431 | 456 |
| **2** | 254 | 549 | 377 | 222 | 262 | 106 | 271 | 301 | 339 |
| **3** | 283 | 595 | 279 | 171 | 184 | 171 | 385 | 246 | 313 |
| **4** | 277 | 443 | 247 | 187 | 163 | 190 | 193 | 197 | 256 |
| **5** | 196 | 517 | 249 | 106 | 197 | 150 | 192 | 176 | 148 |
| **6** | 113 | 581 | 228 | 104 | 141 | 127 | 92 | 106 | 113 |
| **7** | 249 | 491 | 313 | 105 | 158 | 113 | 132 | 119 | 73 |
| **8** | 231 | 606 | 372 | 117 | 149 | 147 | 131 | 106 | 112 |

**Supplementary Table 3**: Primers used in this study.

| Primer name | Sequence |
|---|---|
| pAJM1642_SapI_fw | GGAAAGCTCTTCTTTAGAAAAACTCATCGAGC |
| pAJM1642_SapI_rev | GGAAAGCTCTTCTGCCCAGATAAAATATTTGCTCATG |
| pBBR1_SapI_fw | GGAAAGCTCTTCTTAAGCGGCCACCGGCTGGCTCGCTTCG |
| pBBR1_SapI_rev | GGAAAGCTCTTCTGGCGCATCCTCACGATAATATCC |
| pAJM1642-KanR_NotI_fw | ATAAGAATGCGGCCGCTGTACATCAGAGATTTTGAG |
| pAJM1642-KanR_XbaI_rev | GCTCTAGAGCGGCCACCGGCTGGCTCGC |
| CAT_XbaI_fw | GCTCTAGAGCATCAAATAAAACGAAAGG |
| CAT_NotI_rev | ATAAGAATGCGGCCGCCCAACTTTTGGCGAAAATGAGACG |
| pORM-P12 _SapI_fw | GGAAAGCTCTTCTTAACCCTTCAAATTCAAAGGTAACAAACATGGAAAAACAGACCGAAAATACC |
| pORM-P12 _SapI_rev | GGAAAGCTCTTCTCATGATAGTTACCTTTAAAGTTAAGTGGATTAGGTGCCTTTGATGGTACG |
| P12T2_SapI_fw | GGAAAGCTCTTCTTTAAGCGGCAGGGGCAGG |
| P12T2_SapI_rev | GGAAAGCTCTTCTATGGAAATCGTAAGCAAGCTGACTCTG |
| P12&P19_remove_SapI_fw | GGAAAGCTCTTCTTAACTCGGTACCAAATTCCAGAAAAGAGG |
| P12&P19_remove_SapI_rev | GGAAAGCTCTTCTTTAGGTGCCTTTGATGGTACG |
| KanR_for_pORM_XbaI_fw | GGTGGCCGCTCTAGAGCATCAAATAAAACGAAAGGCTCAGTCGAAAGACTGGGCCTTTCGTTTTA TCTGTTGTTTGTCGGTGAACGCTCTCCTGAGTTTAGAAAAACTCATCGAGC |
| KanR_for_pORM_BsrGI_rev | AATCTCTGATGTACATTGCACAAGATAAAAATATATCATC |
| pORM_P1Y127A_SapI_fw | GGAAAGCTCTTCTCGCTGGCAAGATGGAACG |
| pORM_P1Y127A_SapI_rev | GGAAAGCTCTTCTGCGTCAATTTCAATTTTCTCATCTGATG |
| pOTH_bb_BsaI_fw | GGAAAGGTCTCTGCTAGGATCTCGAGACAGTAAGAC |
| pOTH_bb_BsaI_rev | GGAAAGGTCTCTTTATACCTCTGAATCAATATCAACC |
| p12CO_pOTH_BsaI_fw | GGAAAGGTCTCTATAACCACTTAACTTTAAAGGTAAC |
| p19CO_pOTH_BsaI_rev | GGAAAGGTCTCTTAGCTTACAGAAACAGTTCCAGACG |
| pORFITR_PromChange_AvrII_fw | CACACCTAGGGCCCAAGTTCACTTAAAAAGGAGATCAACAATGAAAGCAATTTTCGTACTGAAAC ATCTTAATCATGCTAAGGAGGTTTTCTAATGTTACGCAGCAGCAACGATGTTACG |
| pORFITR_BG28_AvrII_rev | CACACCTAGGTACATACATTATATCCATGTCAACCTAGTTAATTAAGACGTCTTAAT |
| pORFITR_BG17_AvrII_rev | CACACCTAGGTCATCTGATTATACTCGGCTTTTCGAGTGTTGTCAACTCCGTTAATTAAGACGTCT TAAT |
| GmR_Bpu10I_fw | CTTAATCATGCTAAGGAGGTTTTCTAATGTTACGCAGCAGCAACG |
| GmR_SbfI_rev | AAAGCGATAGCCTGCAGGTTTAGGTGGCGGTACTTGGGTCGATATCAAAGTGC |
| qPCR_dxs_Pputida_fw | GAA GTC AAT GCC GAC ATG CTG G |
| qPCR_dxs_Pputida_rev | GCA CTT TCT TGC TGC CTT CGC |
| qPCR_GmR_fw | AAG TTA GGT GGC TCA AGT ATG G |
| qPCR_GmrR_rev | CGG AAT CCG GCT GAT GTT G |
| qPCR_dxs_Ecoli_fw | CAATTAAAGAGCTGCTCAAACG |
| qPCR_dxs_Ecoli_rev | TTCTTTAGCGTGGTGATAAGC |
